# A screen of chromatin-targeting compounds identifies TAF1 as a novel regulator of HIV latency

**DOI:** 10.1101/2025.05.24.655900

**Authors:** Samuel D. Burgos, Airlie M. Ward, Manickam Ashokkumar, Kimberly P. Enders, Lindsey I. James, David M. Margolis, Edward P. Browne

## Abstract

Antiretroviral therapy (ART) suppresses HIV replication but fails to eliminate the virus due to the persistence of a transcriptionally silent reservoir, which remains the primary barrier to a cure. HIV latency is maintained through chromatin-mediated repression, making epigenetic regulators attractive therapeutic targets. To identify new modulators of latency, we screened a focused library of 84 chromatin-targeting small molecules. This screen identified BAY-299, a bromodomain inhibitor selective for TAF1 and BRD1, as a latency-modulating compound. BAY-299 reactivated HIV expression and enhanced the efficacy of established latency-reversing agents (LRAs), including vorinostat, prostratin, and iBET-151, in cell line models. CRISPR/Cas9-mediated knockout experiments demonstrated that TAF1, but not BRD1, is essential for maintaining HIV latency and that TAF1 depletion selectively increases HIV transcription with minimal effects on host gene expression. Dual knockout of TAF1 and Tat revealed that the reactivation effect is partially Tat dependent. CUT&RUN analysis further showed that TAF1 depletion increases RNA Polymerase II occupancy across the HIV gene body, suggesting enhanced transcriptional elongation. These findings identify TAF1 as a novel regulator of HIV latency and demonstrate the utility of targeted chemical screening to uncover therapeutic vulnerabilities within the latent reservoir.

**Importance:** HIV remains incurable due to the persistence of a transcriptionally silent reservoir in infected cells that is not eliminated by antiretroviral therapy. This transcriptionally silent state, known as latency, is controlled by host cell factors that regulate access to the viral genome. In this study, we identified the host protein TAF1 as a key regulator that maintains HIV in a latent state. Using both genetic and chemical approaches, we demonstrated that reducing TAF1 levels selectively increases HIV gene expression without broadly disrupting host gene transcription. These findings highlight a previously unrecognized mechanism of HIV latency control and identify TAF1 as a potential therapeutic target. Understanding how host chromatin regulators contribute to latency is essential for developing strategies that aim to eliminate the persistent HIV reservoir.

## Introduction

Antiretroviral therapy (ART) has transformed the management of HIV, improving quality of life and extending lifespans for people living with HIV (PLWH) (1–3). However, ART is not curative, as it fails to eliminate a rare but stable reservoir of latently infected cells harboring integrated, transcriptionally silent proviruses (4). These latently infected cells can sporadically reactivate, leading to rebound of viremia if treatment is interrupted (5). Thus, eliminating the latent reservoir remains a key objective in efforts to achieve a functional or sterilizing HIV cure.

To overcome HIV latency, considerable effort has been directed toward developing latency-reversing agents (LRAs) that reactivate latent proviruses, either triggering cytopathic effects or enabling immune-mediated clearance of infected cells (6–13). LRAs such as histone deacetylase (HDAC) inhibitors and inhibitors of apoptosis proteins (IAPs) have shown promise in preclinical models by promoting HIV transcription (14). These compounds function by disrupting proviral silencing. However, most LRAs exhibit limited potency and breadth of reactivation, underscoring the need for a deeper understanding of latency mechanisms to inform alternative or combinatorial therapeutic strategies.

The persistence of the HIV reservoir is driven by host chromatin regulation, particularly covalent histone modifications, such as acetylation and methylation, that shape proviral chromatin structure. Histones H3 and H4 possess modifiable tails targeted by protein complexes that read, write, or erase these marks (15, 16). Once integrated, HIV is governed by the same chromatin-based mechanisms that regulate host transcription, including histone modifications and nucleosome remodeling (17–19). These processes restrict access to the transcriptional machinery, maintaining latency and suppressing reactivation. Given the complexity of these pathways, chromatin regulation offers multiple potential therapeutic targets. Although some LRAs disrupt silencing mechanisms, most fail to reactivate the bulk of the latent reservoir (20, 21), highlighting the need to identify additional compounds that engage distinct regulatory pathways and possible combinations with current LRAs.

To identify novel chromatin-associated factors that regulate HIV latency, we screened a library of 84 small molecules targeting enzymes involved in writing or erasing histone modifications, including bromodomain proteins (BRDs), lysine demethylases (KDMs), histone acetyltransferases (HATs), deacetylases (HDACs), methyltransferases (HMTs), and methyl-lysine readers (Kme readers). The screen identified latency-modulating compounds across several epigenetic classes, with particularly strong activity among BRD and HDAC inhibitors. One notable hit was BAY-299, a bromodomain antagonist selective for TAF1 and BRD1 (22), which enhanced HIV expression and sensitized latently infected cells to canonical LRAs. CRISPR/Cas9 knockout experiments confirmed that TAF1, not BRD1, is required to maintain HIV latency. These findings demonstrate the power of targeted chemical screens to reveal novel regulatory pathways and therapeutic vulnerabilities within the latent HIV reservoir.

## Results

### A chemical screen identifies novel compounds affecting HIV latency

To identify novel compounds that affect HIV latency, we screened an 84-compound library targeting proteins that add or remove post-translational histone modifications. The screen was conducted in Jurkat cells, a human CD4⁺ T cell line, infected with the HIV-GKO strain (23). HIV-GKO is a dual-color reporter virus that maintains expression of most viral genes but includes a nonfunctional env gene and two fluorophores: eGFP under the HIV-1 5’ LTR promoter and mKO under the EF1α promoter within the nef gene (**Figure 1A**). This virus thus enables quantification of both active (eGFP+/mKO+) and latent (eGFP-/mKO+) infection within an infected population of cells (**Figure 1B**).

**Figure 1.**
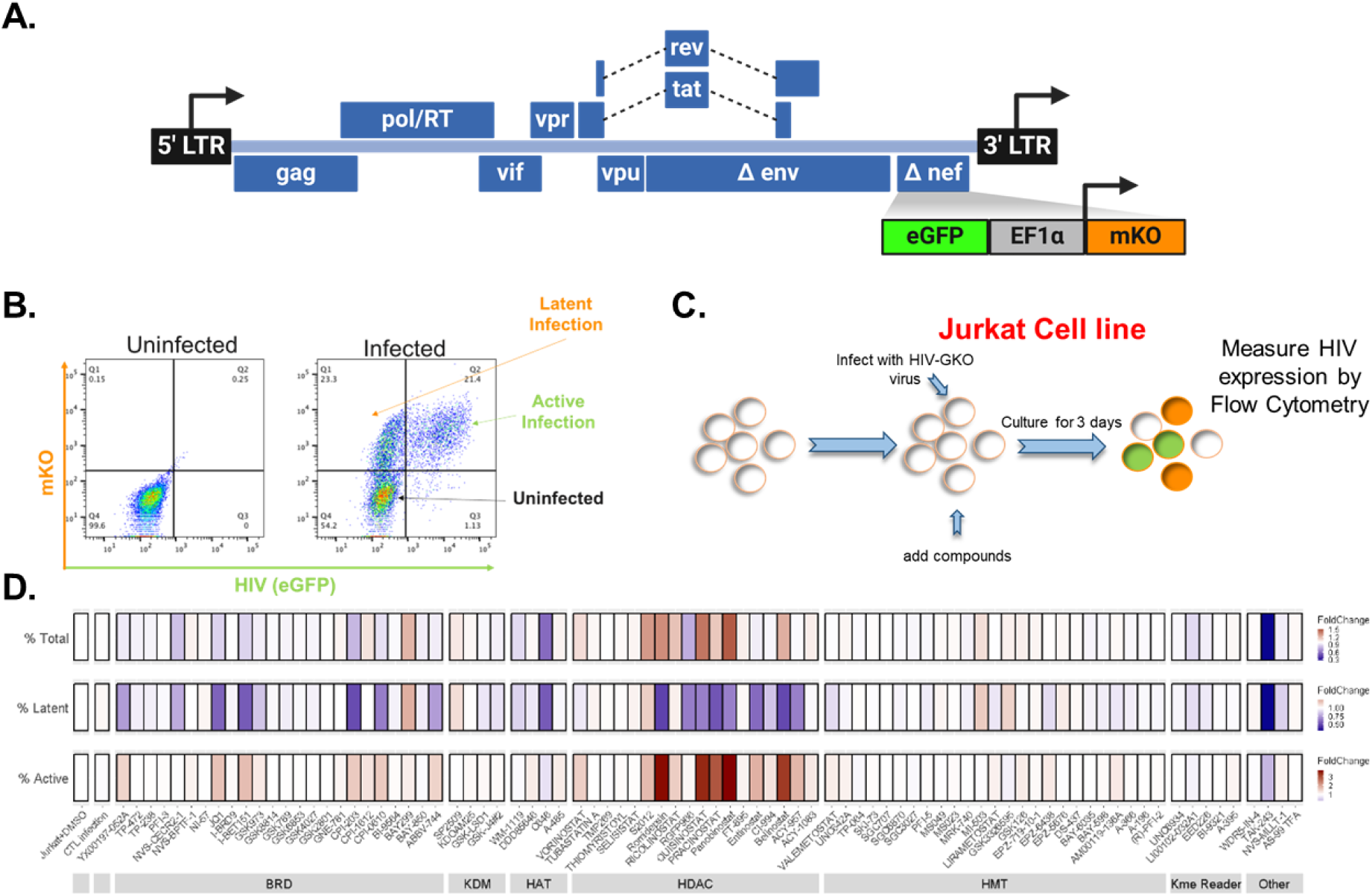
Chemical screen identifies novel chemical compounds affecting HIV latency. **A**. The HIV-GKO gene map illustrating the expressed viral genes and the location and identity of the fluorophore cassettes, adapted from E. Battivelli *et al* (23). **B**. Example of flow cytometry of HIV-GKO infected Jurkat cells, highlighting latent and active infections based on the viral promoter driving GFP expression and the constitutive promoter EF1α driving mKO expression. **C**. Schematic representation of the small molecule compound screening in Jurkat cell line model of HIV latency. **D**. A multivariable heatmap illustrates changes relative to the DMSO control for total infection percentage, latent infection percentage, and active infection percentage. Data shown are the average of four replicates.

HIV-GKO-infected Jurkat cells were treated with each compound at 1 µM in a 96-well plate format to enable high-throughput screening **(Figure 1C**). After three days, cells were analyzed by flow cytometry to assess changes in infection profiles. Live cells were gated using a viability dye, then assessed for mKO expression to determine total infection (% Total Infection). mKO-positive cells were further gated based on eGFP to define latent (eGFP⁻/mKO⁺) and active (eGFP⁺/mKO⁺) infections. Fold changes for each population were calculated relative to DMSO controls (**Figure 1D**). None of the compounds screened significantly altered total infection (all FDR > 0.05). An increase in the percentage of latently infected cells was observed for 25 compounds under these conditions (FDR < 0.05): nine bromodomain antagonists (ABBV-744, BAY-299, CPI-0610, CPI-203, GSK973, I-BET151, JQ1, NVS-CECR2-1, YX00197-052A1), the KDM inhibitor SP2509, two HAT inhibitors (A-485, C646), eleven HDAC inhibitors (ACY-957, belinostat (S1085), CI-994, panobinostat (S1030), entinostat (S1053), pracinostat (S2012), quisinostat, ricolinostat, RGFP966, romidepsin (S3020), vorinostat), the PRMT5 inhibitor GSK3326595, and the ubiquitin-activating enzyme inhibitor TAK-243. Conversely, nine compounds—ABBV-744, CPI-0610, CPI-203, GSK973, I-BET151, JQ1, NVS-CECR2-1, PFI-3, and C646—produced a significant decrease in the latent cell population (**Table 1**). Finally, 23 compounds significantly increased the percentage of actively infected cells (FDR < 0.05), including ABBV-744, BAY-299, CPI-0610, CPI-1612, CPI-203, GSK973, I-BET151, JQ1, A-485, belinostat, CI-994, panobinostat, entinostat, pracinostat, quisinostat, ricolinostat, RGFP966, romidepsin, S2012, vorinostat, EPZ-6438, FL00040-097B1, and TAK-243, whereas only TAK-243 significantly reduced active infection (**Table 1**). The results of this screen approach demonstrate the approaches’ capability to identify novel small molecules that influence HIV expression and latency.

**Table 1:** List of compounds with an effect on an HIV infected population. The list of compounds that modulated abundance of % total infected, % latently infected or % actively infected is shown. Data is listed as fold change (FC) relative to control (DMSO) exposed cells. Arrow indicates direction of change.

| Compound | Class | Total FC | Latent FC | Active FC |
| --- | --- | --- | --- | --- |
| ABBV-744 | Bromodomain | 1.3 | 0.53 ↓ | 1.47 ↑ |
| BAY-299 | Bromodomain | 1.6 | 0.87 ↑ | 1.25 ↑ |
| CPI-0610 | Bromodomain | 1.2 | 0.47 ↓ | 1.47 ↑ |
| CPI-1612 | Bromodomain | 1.5 | 0.7 | 1.25 ↑ |
| CPI-203 | Bromodomain | 1.1 | 0.33 ↓ | 1.47 ↑ |
| GSK973 | Bromodomain | 1.3 | 0.60 ↓ | 1.25 ↑ |
| I-BET151 | Bromodomain | 1.2 | 0.40 ↓ | 1.75 ↑ |
| JQ1 | Bromodomain | 1.3 | 0.40 ↓ | 1.75 ↑ |
| NVS-CECR2-1 | Bromodomain | 1.1 | 0.53 ↓ | 1.0 |
| YX00197-052A1/<br>GSK525762/ I-BET762 | Bromodomain | 1.3 | 0.5 | 1.5 |
| SP2509 | Demethylase | 1.5 | 0.8 | 1.0 |
| A-485 | HAT | 1.4 | 0.4 | 1.25 ↑ |
| C646 | HAT | 0.82 ↓ | 0.40 ↓ | 0.75 ↓ |
| ACY-957 | HDAC | 1.75 ↑ | 0.5 | 1.75 ↑ |
| Belinostat | HDAC | 1.6 | 0.33 ↓ | 3.50 ↑ |
| CI-994 | HDAC | 1.3 | 0.60 ↓ | 1.25 ↑ |
| Panobinostat | HDAC | 2.00 ↑ | 0.5 | 3.00 ↑ |
| Entinostat | HDAC | 1.3 | 0.4 | 2.00 ↑ |
| Pracinostat | HDAC | 1.6 | 0.8 | 2.00 ↑ |
| Quisinostat | HDAC | 1.9 | 0.5 | 3.00 ↑ |
| Ricolinostat | HDAC | 1.6 | 0.7 | 1.50 ↑ |
| RGFP966 | HDAC | 1.5 | 0.5 | 1.0 |
| Romidepsin | HDAC | 1.8 | 0.33 ↓ | 3.75 ↑ |
| Vorinostat | HDAC | 1.5 | 0.7 | 1.50 ↑ |
| GSK3326595 | HMT | 1.4 | 0.8 | 0.75 ↓ |
| EPZ-6438 | HMT | 1.4 | 0.7 | 1.25 ↑ |
| UNC1999A | HMT | 1.4 | 0.7 | 1.25 ↑ |
| GSK126 | HMT | 1.4 | 0.7 | 1.25 ↑ |
| SGC707 | HMT | 1.4 | 0.7 | 1.25 ↑ |
| MI-503 | HMT | 1.5 | 0.8 | 1.0 |
| TAK-243 | Unclassified | 0.36 ↓ | 0.20 ↓ | 0.36 ↓ |

### BAY-299 affects latency reversal in a cell line model of HIV latency

In addition to known LRAs, the screen identified novel compounds, including several from classes not previously studied in the context of HIV. We selected four for further analysis: two that increased total infection, BAY-299 (a TAF1/BRD1 antagonist) and MI-503 (a Menin-MLL inhibitor), and two that elevated the ratio of active to latent infection, PFI-3 (a SMARCA2/4 antagonist) and Tubastatin A (an HDAC6 inhibitor). To validate their activity, we performed dose-response analyses in HIV-GKO-infected Jurkat cells across concentrations from 1 nM to 10 µM, measuring infection profiles by flow cytometry at 3 days post-infection. BAY-299 and PFI-3 both increased total infection at intermediate doses (10–100 nM) (**Figure 2A**). At 1 µM, BAY-299 also modestly increased latent infection, consistent with initial screen results, while other compounds showed no significant change from DMSO controls. Notably, active infection (eGFP⁺/mKO⁺) was significantly elevated by BAY-299 and PFI-3 at 10–100 nM. BAY-299 induced the strongest activation within this range but showed reduced effect at higher doses (1–10 µM). Similarly, PFI-3 peaked between 10–100 nM. These results suggest that while 1 µM was used in the initial screen, both compounds exhibit peak activity around 100 nM. BAY-299 thus shows a concentration-dependent dual effect enhancing activation at lower doses while modestly increasing latency at higher concentrations.

**Figure 2.**
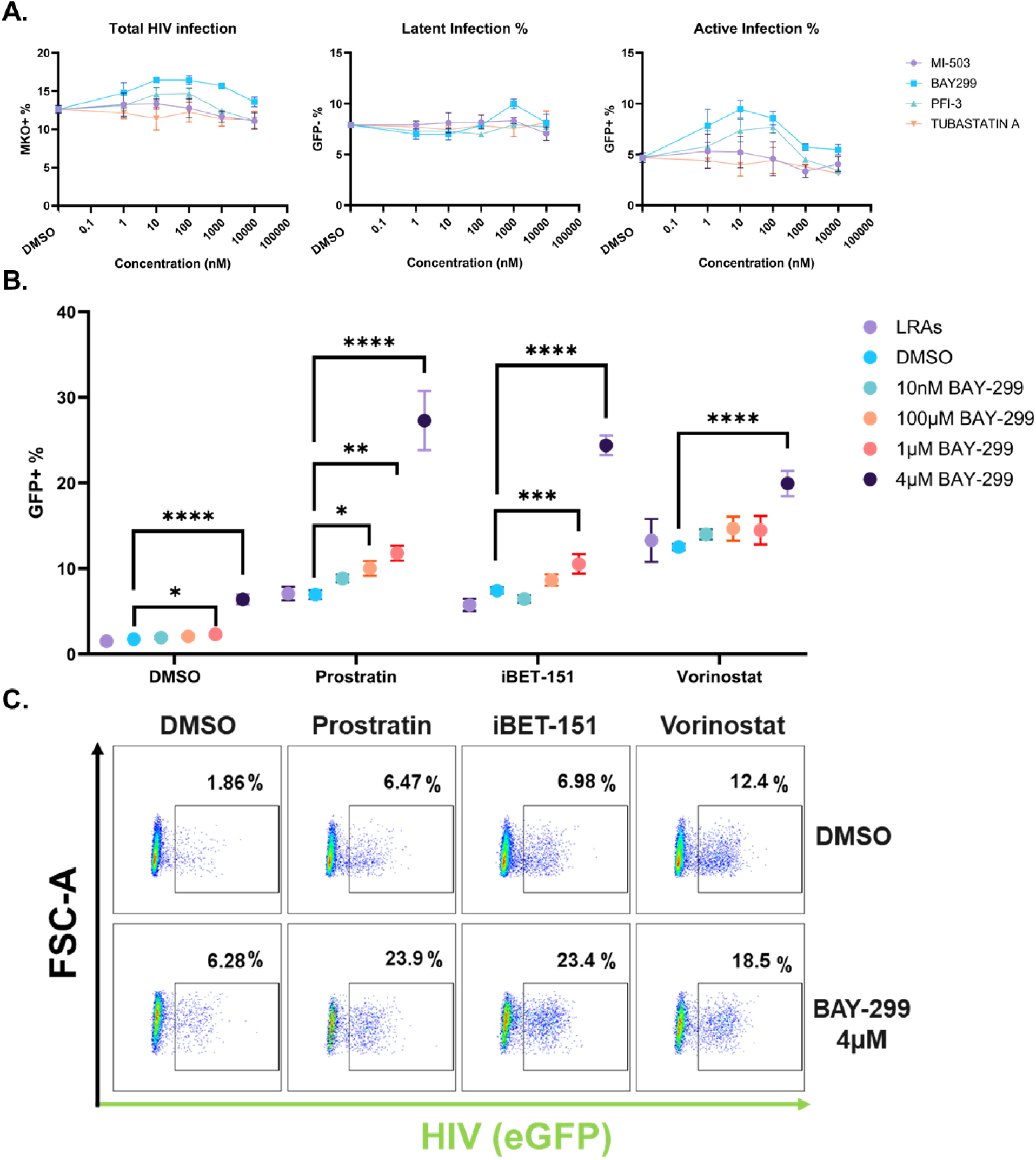
BAY-299 affects HIV gene expression and enhances latency reversal. **A.** Dose-response curves for selected compounds: MI-503, BAY-299, PFI-3, and Tubastatin A, showing the effects on total HIV infection (mKO+), latent infection (mKO+, GFP-), and active infection (mKO+, GFP+) percentages. DMSO control is plotted as a single dot on the plot. Datapoints represent the average of technical triplicates. **B.** Dose-response curve for BAY-299 in 2D10 cells ranging from 10 nM to 4 µM, with and without the addition of LRAs prostratin (125 nM), iBET-151 (125 nM), and vorinostat (250 nM). Measurements are an average of four technical replicates. p-value: ****p<0.0001, ***p<0.001, **p<0.01, *p<0.05. **C.** Flow cytometry showing the changes in GFP expression of 2D10 cells when treated with LRAs alone and LRAs in conjunction with BAY-299 at 4 µM.

We next tested BAY-299 in 2D10 cells, a well-established model of HIV latency containing a quiescent provirus that encodes Tat, Env, Vpu, Rev, and eGFP under the 5′ LTR promoter (24). Cells were treated with BAY-299 at concentrations from 10 nM to 4 µM for 24 hours. Unlike the HIV-GKO model, where peak activation occurred at 10– 100 nM, BAY-299 induced modest reactivation in 2D10 cells only at 1 µM and 4 µM (**Figure 2B**). We also tested whether BAY-299 could enhance the activity of other LRAs: prostratin (125 nM), iBET-151 (125 nM), and vorinostat (250 nM), by co-treating 2D10 cells and measuring HIV expression via eGFP at 24 hours post-stimulation. Notably, combining BAY-299 at 1 µM or 4 µM with any of these LRAs significantly increased GFP expression compared to LRA treatment alone (**Figure 2B and 2C**). This enhancement was strongest in combination with prostratin or iBET-151, producing a threefold increase relative to DMSO controls. These results show that BAY-299 not only has modest latency-reversing activity in 2D10 cells but can significantly potentiate the effects of diverse LRAs.

### TAF1 is required for HIV latency in cell line models

BAY-299 was initially developed as a selective antagonist for Bromodomain and PHD Finger-Containing Protein 2 (BRPF2/BRD1) (22). Although it lacks affinity for the related proteins BRPF1 and BRPF3, the same study reported potent binding to the second bromodomain of TAF1. We hypothesized that BAY-299 affects HIV latency by targeting either BRD1 or TAF1. To test this, we nucleofected CRISPR-Cas9 ribonucleoprotein (RNP) complexes targeting TAF1, BRD1, or a non-targeting control into 2D10 cells. Western blot confirmed robust protein depletion by 5 days post-nucleofection (DPN) (**Figure 3A**). HIV expression was assessed by measuring eGFP fluorescence via flow cytometry at 5 and 7 DPN. Notably, TAF1 depletion, but not BRD1 depletion, significantly increased viral GFP expression (**Figure 3B**), suggesting that TAF1 is important for maintaining HIV latency in 2D10 cells.

**Figure 3.**
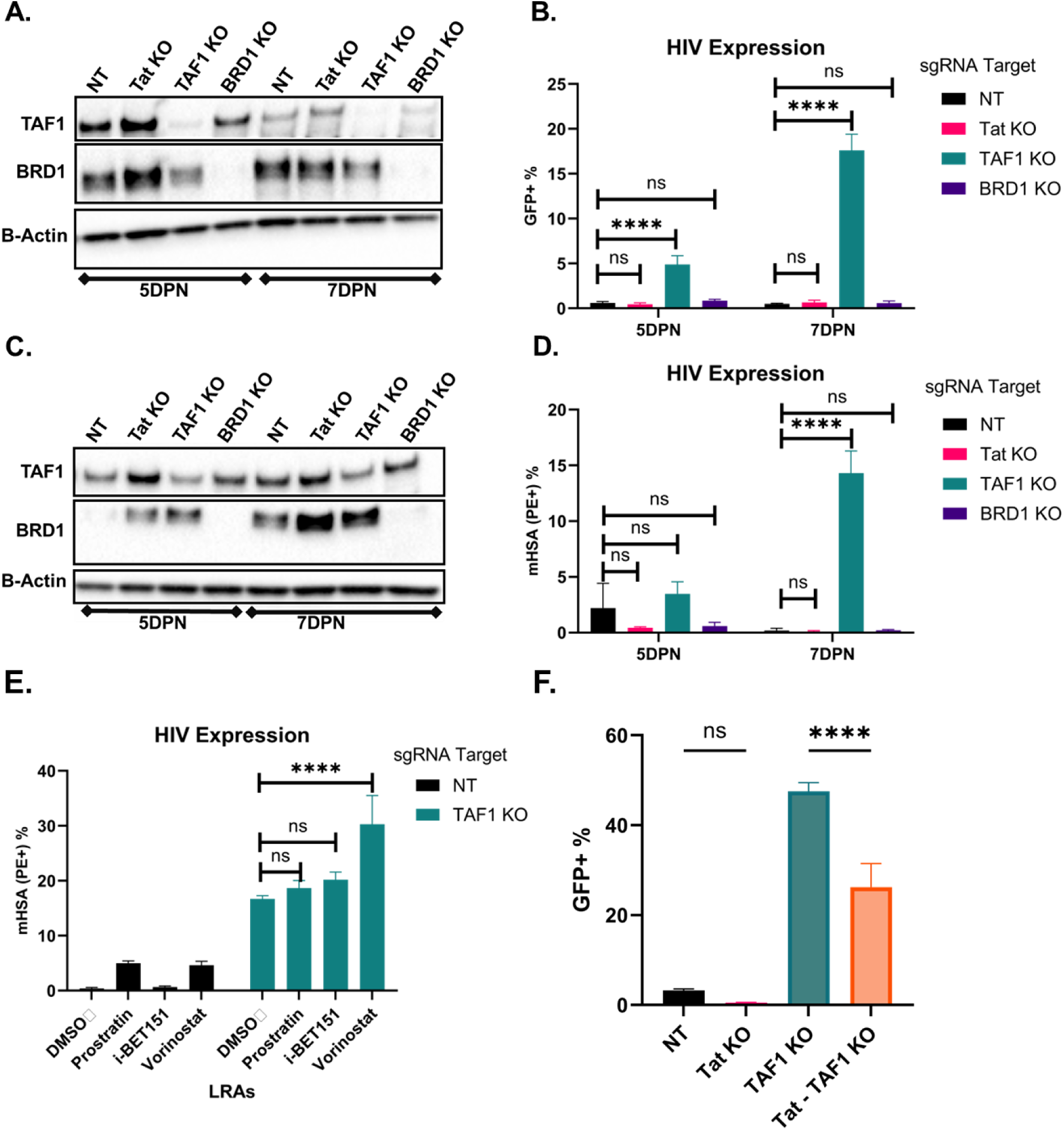
TAF1 depletion leads to Tat-dependent HIV reactivation. **A.** Western blot of 2D10 cells following CRISPR/Cas9 nucleofection targeting TAF1 and BRD1 at 5 days and 7 days post-nucleofection (DPN). Non-targeting gRNA (NT) represents a negative control. **B.** Bar graph showing the changes in average eGFP expression determined by flow cytometry from three replicates in 2D10 cells, comparing the effects of TAF1 and BRD1 depletion. p-value: ****p<0.0001. **C.** Western blot of CRISPR/Cas9-depleted N6 cells targeting Tat, TAF1 and BRD1 at 5 and 7 DPN. **D.** Bar graph showing the changes in average eGFP expression determined by flow cytometry from three replicates of N6 cells, comparing the effects of Tat, TAF1 and BRD1 depletion. **E.** Bar graph showing the effect of 24h treatment of different LRAs (prostratin 250 nM, iBET151 250 nM, vorinostat 500 nM) on the expression of HIV (measured by mHSA/PE+) in the N6 cell line with or without TAF1 depletion at 7DPN. p-value: ****p<0.0001. **F.** Bar graph showing the effect of TAF1 knockout, Tat knockout, and TAF1/Tat double knockout on the percentage of eGFP-positive cells in the 2D10 cell line at 7DPN. p-value: ****p<0.0001.

To further validate this result, we tested TAF1 and BRD1 depletion in a second latency model, the N6 cell line. N6 cells harbor a full-length HIV provirus and encode mouse heat-stable antigen (mHSA) in the Nef open reading frame, allowing quantification of HIV expression via flow cytometry (25). Compared to 2D10 cells, TAF1 depletion in N6 cells was incomplete, while BRD1 knockdown was substantial by 5 and 7 DPN (**Figure 3C**). Despite this, partial TAF1 depletion led to a significant increase in HIV expression, as measured by mHSA staining at day 7 (**Figure 3D**). These data support a consistent role for TAF1, but not BRD1 in repressing HIV across multiple latency models.

We next examined whether TAF1 or BRD1 depletion could enhance HIV reactivation in response to latency-reversing agents (LRAs). N6 cells depleted for TAF1 or BRD1 were treated with suboptimal doses of vorinostat, prostratin, or iBET-151. Among these, only vorinostat exhibited an additive effect with TAF1 depletion, while the others showed no additional increase in HIV expression (**Figure 3E**). This suggests that TAF1 loss can independently reverse latency and selectively enhance the activity of certain LRAs.

To further investigate the mechanism by which TAF1 depletion increases HIV expression, we performed dual knockouts of TAF1 and Tat, the HIV-encoded transcriptional activator. Prior studies have shown that even without multiple TAFs, including TAF1, components of the pre-initiation complex can still bind the HIV promoter, with Tat thought to compensate (26). In 2D10 cells, we nucleofected RNPs targeting TAF1, Tat, or both. Consistent with prior results, TAF1 depletion alone elevated HIV expression (**Figure 3F**). Tat depletion alone had no effect, likely due to low basal activity in latently infected cells. However, combined depletion of TAF1 and Tat significantly reduced the increase observed with TAF1 knockout alone, though levels remained above those of the NT or Tat-only groups. These results suggest that Tat contributes to the transcriptional enhancement observed following TAF1 depletion.

### TAF1 depletion in 2D10 cells results in the preferential upregulation of HIV transcription

TAF1 has been reported to facilitate transcriptional initiation by recruiting the TFIID complex to acetylated histones at gene promoters (27, 28). We hypothesized that TAF1 depletion might directly affect HIV transcription or alter expression of host genes that regulate latency. To test this, we depleted TAF1 in 2D10 cells using CRISPR-Cas9 nucleofection and confirmed increased viral GFP expression by flow cytometry (**Figure 4A**). RNA from TAF1-depleted and non-targeting control cells was analyzed by bulk RNA-sequencing. Differential expression analysis using DESeq2 identified 163 significantly upregulated and 354 significantly downregulated genes (padj < 0.05) in TAF1-depleted cells (**Figure 4B**). Notably, HIV was the most significantly upregulated transcript, suggesting that TAF1 plays a relatively selective role in repressing HIV expression in this model. Pathway enrichment analysis revealed no dominant functional pathways among DEGs, indicating that TAF1 depletion drives robust HIV reactivation with only modest effects on host gene transcription.

**Figure 4.**
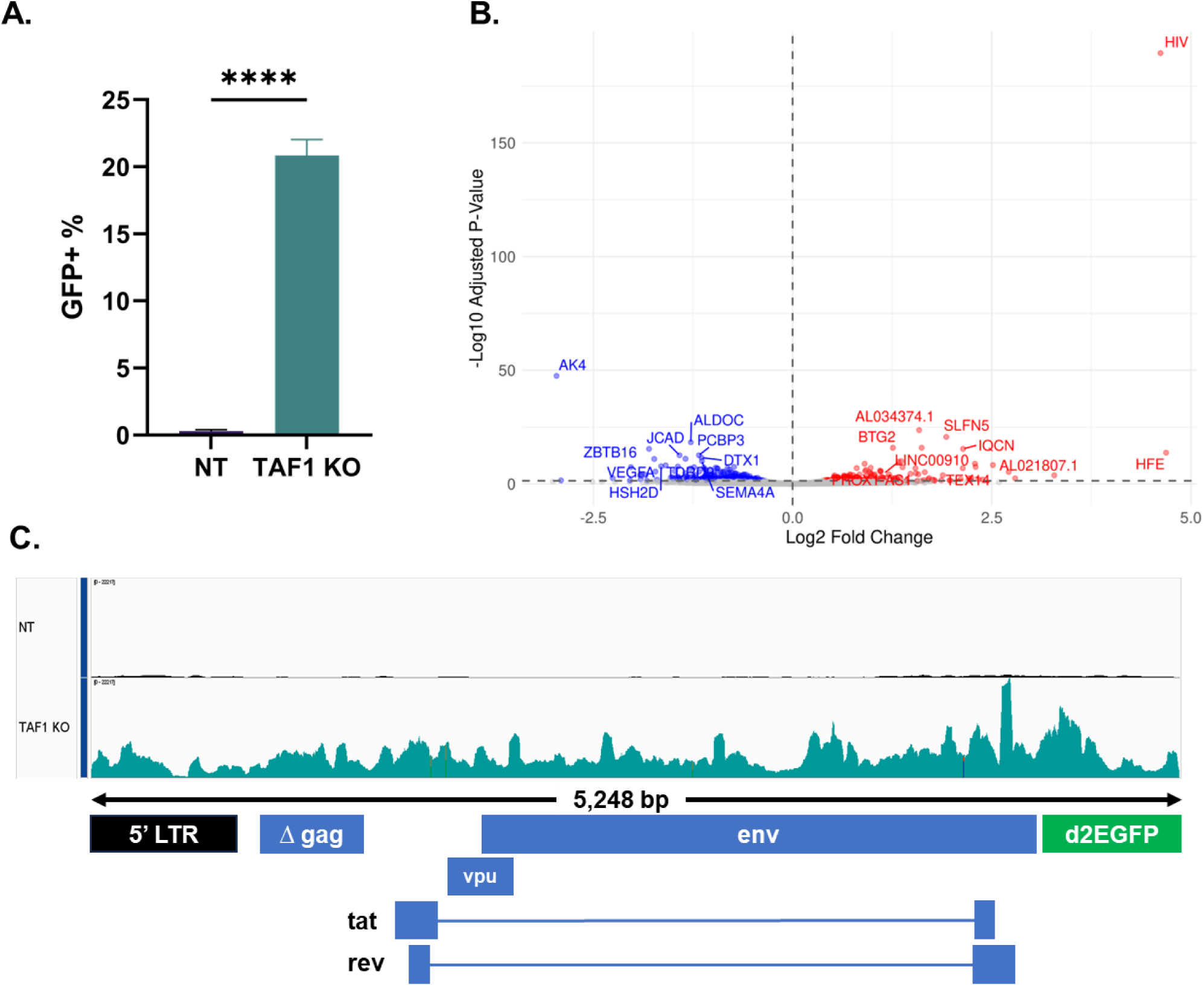
TAF1 depletion in 2D10 cells results in a preferential increase in HIV transcripts. **A.** Bar graph showing that TAF1 depletion in 2D10 cells leads to a significant increase in eGFP+ cells at 7 days post nucleofection, indicating elevated HIV expression in the same experimental context used for transcriptomic profiling. p-value: ****p<0.0001. **B.** Volcano plot highlighting significantly upregulated (red) and downregulated (blue) genes in TAF1-depleted cells versus non-targeting controls. Colored points represent significant genes with adjusted p-values (log p-adjusted) less than 0.05, while grey points represent that are non-significant. **C.** RNAseq coverage tracks of the HIV genome in 2D10 cells, comparing non-targeting CRISPR/Cas9 treated cells and TAF1-depleted cells.

### TAF1 depletion regulates RNAPol II abundance across the HIV gene body

To investigate how TAF1 depletion enhances HIV transcription in 2D10 cells, we performed Cleavage Under Targets and Release Using Nuclease (CUT&RUN) to examine activating histone marks and RNAPol II at the HIV provirus. Cells were nucleofected with CRISPR/Cas9 RNPs targeting TAF1 or a non-targeting control (NT) (**Figure 5A**). At 7 days post-nucleofection, HIV reactivation was confirmed by increased eGFP expression (**Figure 5B**). CUT&RUN was conducted using antibodies against H3K4me3 and H3K9ac (activating histone marks), RNAPol II, and IgG control. As expected, we observed enrichment of H3K4me3, H3K9ac, and RNAPol II at the HIV LTR near the transcription start site (TSS), consistent with prior reports (29). However, TAF1-depleted cells showed no substantial change in the abundance of these marks at the LTR, despite elevated HIV expression (**Figure 5C, 5D**). This indicates that TAF1 represses HIV through a mechanism independent of increased histone activation marks or RNAPol II recruitment at the promoter. In contrast, we observed a notable increase in RNAPol II occupancy across the HIV gene body. Specifically, there was an ∼2-fold increase in RNAPol II signal between base pairs 1,000 and 5,248, spanning the region downstream of the LTR through the end of the provirus in the 2D10 model (**Figure 5D)**. While RNAPol II levels at the LTR remained constant, its gene body abundance increased, suggesting that TAF1 depletion enhances transcriptional elongation or RNAPol II processivity.

**Figure 5.**
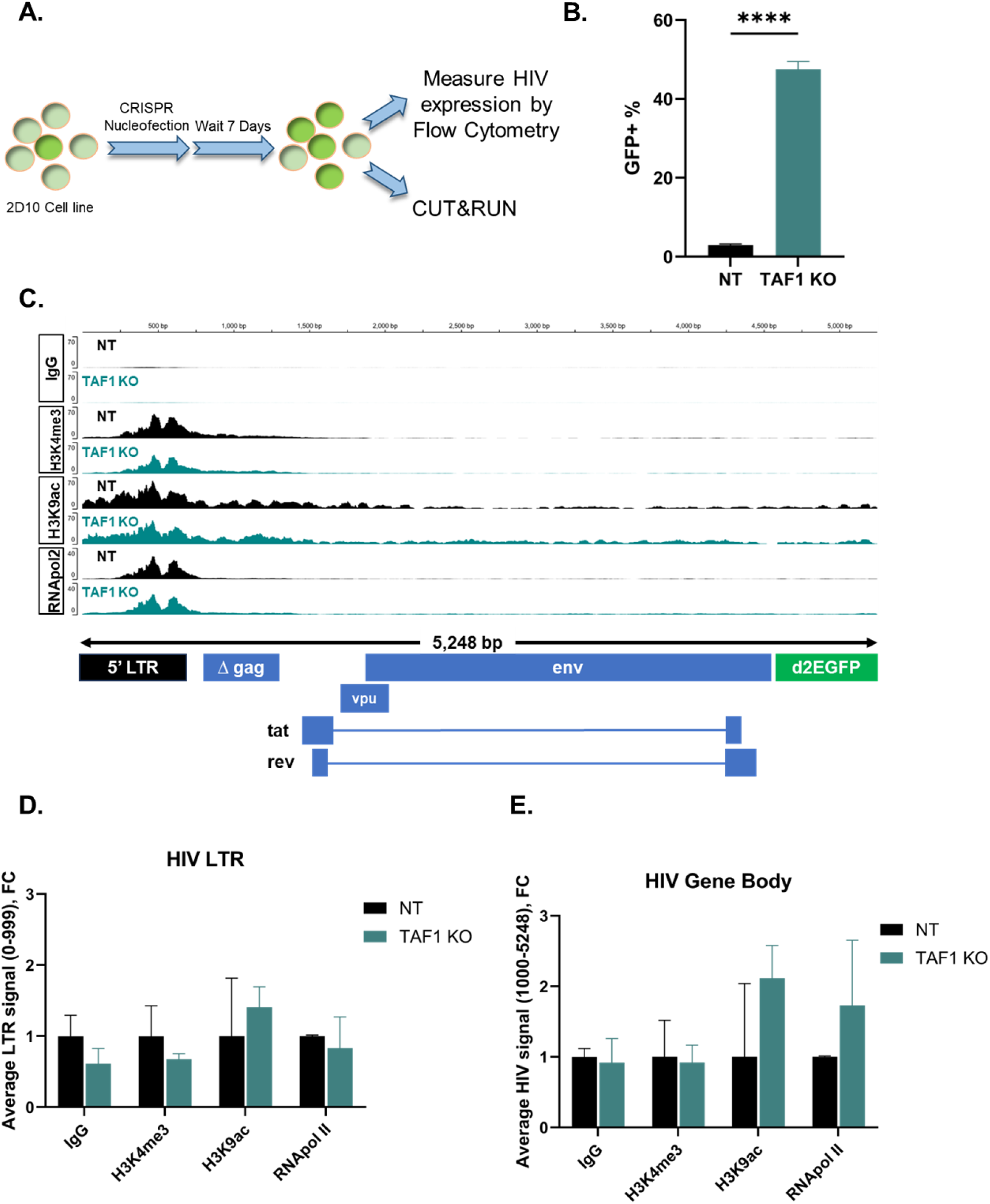
TAF1 depletion in 2D10 cells increases RNAPol II abundance within the HIV gene body. **A.** Schematic illustration of CUT&RUN experimental design. 2D10 cells were nucleofected with either NT or TAF1-targeting ribonucleoprotein (RNP) complexes. At 7 days post nucleofection (DPN), cells were analyzed for GFP expression by flow cytometry and processed for CUT&RUN profiling. **B.** Bar graph showing increase in eGFP+ cells following TAF1 depletion in 2D10 cells at 7 DPN, p-value: ****p<0.0001. **C.** CUT&RUN sequence coverage of the HIV genome. Tracks correspond to coverage for either IgG (negative control), H3K4me3, H3K9ac, or RNAPol II for NT or TAF1 depleted 2D10 cells. **D.** Bar graph showing fold change read counts for IgG, H3K9ac, H3K4me3, and RNAPol II (Rbp1) immunoprecipitations within the HIV LTR region (nts 0-1000). **E.** Bar graph showing fold change read counts for IgG, H3K9ac, H3K4me3, and RNAPol II (Rbp1) immunoprecipitations across the HIV gene body (nts1000 to 5248).

## Discussion

The persistence of latently infected cells remains the primary barrier to achieving an HIV cure, as this reservoir evades immune surveillance and resists ART (30, 31). Despite significant advancements in LRAs, there is still limited efficacy in fully reactivating the latent reservoir, highlighting the need for novel therapeutic approaches (32). LRAs such as panobinostat and vorinostat (HDACi), JQ1 and iBET151 (BRD4i) and AZD5582 (Inhibitor of Apoptosis Proteins) are well-studied compounds that have demonstrated the ability to increase HIV gene expression via different mechanisms. However, individually, they have not been able to broadly reactivate the latent reservoir (33).

In this study we found that many of the compounds that increased the percentage of actively expressing HIV cells are members of the HDAC inhibitor family or bromodomain 4 (BRD4) antagonist that interfere with the binding of BRD4 to acetylated histones. This observation highlights the central importance of histone acetylation in HIV latency, consistent with previous reports (7, 10, 20, 34, 35). Several of these compounds have previously been identified as LRAs, including the HDACis romidepsin, panobinostat, and vorinostat, as well as the BRDis iBET-151 and JQ1. This observation thus gives us confidence in our screen by validating that we were able to identify known LRAs. Compounds from the other categories in the library did not affect the percent active population, except for TAK-243, an inhibitor of ubiquitin activating enzyme (UAE), which reduced the percent active population, highlighting a different avenue to affect HIV transcription.

From the overall set of biologically active compounds identified by this screen, we selected BAY-299 for further analysis. Subsequent analysis confirmed BAY-299’s ability to increase the percent total infection, percent latent, and percent active infection. Notably, when used in combination with different LRAs such as prostratin, iBET-151, and vorinostat, 1 μM or 4 μM BAY-299 further enhanced HIV transcription, demonstrating a possible combinatorial effect on HIV transcription. BAY-299 is a bromodomain antagonist specifically designed to differentiate between bromodomain paralogs, with selectivity for BRD1 and TAF1 (22). Given that this compound influenced all measured parameters related to changes in HIV latency and transcription, we further investigated the possible mechanism by which BAY-299 affects HIV by selective depletion of its targets BRD1 or TAF1. Notably, depletion of TAF1 and not BRD1 caused a robust increase in HIV transcription in two cell lines that were latently infected with HIV (2D10 cells and N6 cells). Additionally, the changes in gene expression caused by TAF1 depletion appear to be relatively selective for HIV transcription, with minimal impact on host gene expression. This observation is particularly intriguing because TAF1 is a subunit of the TFIID complex, which plays a an important role in transcriptional initiation as the rate-limiting step for RNA polymerase II binding to DNA (27). Given TAF1s well known role in facilitating transcriptional initiation, it is curious that TAF1 exhibits a specific HIV-repressing activity. Notably, previous *in vitro* studies have demonstrated that HIV transcription can occur in the absence of TAF subunits within the TFIID complex, including TAF1, suggesting the existence of compensatory mechanisms that enable transcription despite their depletion (26). It has been proposed that HIV maintains transcription under these conditions through a compensatory mechanism involving Tat, an essential viral protein for HIV transcription (26). This hypothesis aligns with our RNA-seq data that demonstrate a specific increase in HIV transcription in the absence of TAF1.

Previous work has shown that TAF7 contains a domain that binds TAF1, enabling formation of the TFIID pre-initiation complex and simultaneously inhibiting the acetyltransferase activity of TAF1. This interaction is thought to function as a transcriptional checkpoint regulated by TAF7 (36). It was also demonstrated that HIV-1 Tat, the HIV-encoded transcriptional activator, contains a structurally similar domain to TAF7, allowing it to bind TAF1 in a comparable manner. This interaction has been proposed to downregulate the expression of TAF1-dependent host genes, such as MHC class I, potentially contributing to immune evasion by reducing antigen presentation (37).

Based on these findings, we hypothesize that in our HIV cell line models, depletion of TAF1 reduces the availability of TAF1 for Tat binding. As a result, more Tat may be available to drive transcription at the HIV promoter, leading to the observed increase in HIV gene expression. This is supported by the robust increase in HIV-1 mRNA levels and increase in eGFP expression in TAF1-depleted cells. Additionally, our CUT&RUN data reveals elevated RNA polymerase II occupancy across the HIV gene body in the absence of TAF1, consistent with enhanced transcriptional activity. Taken together, these data are consistent with a model in which TAF1 depletion relieves Tat sequestration, allowing greater Tat availability to promote HIV transcription.

Our results from this screen should be considered in light of several limitations and caveats. While this screen was effective in identifying novel compounds that influence HIV latency, it was performed using immortalized cell lines as the model for measuring changes in HIV transcription, which exhibit several differences from primary CD4 T cells, the primary location of the clinical reservoir (38, 39). Consequently, it is unknown if the findings from this screen will fully translate to primary cells or in vivo models of HIV latency. The library used in this screen consisted of 84 distinct small molecules that target various chromatin functions critical to transcription. Due to the variability in efficacy, selectivity profiles, and toxicity of these compounds, the optimal dose for each compound likely differs. To standardize the approach, however, we selected a single concentration of 1 µM, a dose low enough to minimize toxicity yet sufficient to be effective in most cases, recognizing that this dose is likely not optimal for all compounds tested. Nevertheless, at this concentration, we observed notable effects in response to several compounds. Future screening could potentially examine the compound set across multiple concentrations in order to more thoroughly evaluate the biological activity of the molecules. Another variable to consider is the timing of compound treatment. In this screen, compounds were added on the same day as infection to capture changes in total infection, latent infection, and active infection. However, as the HIV integration complex takes 24–48 hours to fully integrate into the host genome, the timing of compound addition could influence the results.

Despite these limitations, this screen serves as a robust template for identifying novel epigenetic mechanisms and pathways that can influence HIV transcription. Through this approach, we identified BAY-299, a compound that affects HIV latency by targeting TAF1 and modulating HIV transcription.

## Methods

### Cell line model of HIV latency

The 2D10 and Jurkat-N6 latently infected cell lines (40), along with parental Jurkat cells, were maintained in RPMI-1640 medium (Gibco, Thermo Fisher), supplemented with 10% fetal bovine serum (FBS), 2 mM L-glutamine, sodium pyruvate, 100 U/mL penicillin-streptomycin, and 10 mM HEPES. Cells were cultured at 37 °C with 5% CO₂. Latently infected Jurkat cells were generated via infection with the HIV-GKO virus. HEK293T cells (ATCC, CRL-11268), used for transfection and viral production, were maintained in DMEM (Gibco) with 10% FBS and penicillin-streptomycin.

### Virus stocks

HIV-GKO viral stocks were produced by co-transfecting HEK293T cells (ATCC CRL-3216) with the HIV-GKO plasmid (Addgene #112234) and the VSV-G envelope plasmid pMD2.G (Addgene #12259) using Mirus-LT1 reagent (Mirus Bio, MIR2305) at a 1:4 VSV-G:HIV-GKO plasmid ratio. Transfected cells were cultured in DMEM (Gibco, 11995-065) with 10% FBS. At 48 hours post-transfection, supernatants were centrifuged at 600 × g for 5 min and filtered through 0.45 μm membranes to remove debris, then aliquoted and stored at −80 °C. Viral titers were determined by infecting Jurkat T cells (ATCC TIB-152), optimizing MOI to avoid multiple infections. Jurkat cells were maintained in complete RPMI for up to three days post-infection.

### Small molecule inhibitor screen

A focused epigenetic inhibitor library consisting of 84 small molecules (assembled by the James Lab, UNC-CH) was initially dispensed at 10 mM in a 384-well format. Compounds were transferred to 96-well plates and diluted to 1 mM in DMSO, then further diluted to 200 µM in RPMI (1:5) to maintain final DMSO concentrations ≤0.1%. For screening, 1 µL of each 200 µM compound was added to V-bottom 96-well plates containing 30,000–50,000 HIV-GKO-infected Jurkat cells per well, 2 hours post-infection, in a final volume of 200 µL. The layout was repeated across four technical replicate plates. DMSO vehicle controls were prepared using the same dilution scheme. After three days of concurrent infection and treatment, plates were centrifuged at 300 × g for 5 minutes and cells were processed for flow cytometry.

### Flow Cytometry

Viral gene expression was assessed by flow cytometry using mKO (HIV-GKO), eGFP (2D10), or the surface marker mouse Heat Stable Antigen (mHSA) (N6; BD Biosciences, 553262), depending on the model system. Cell viability was determined using Zombie Violet dye (BioLegend, 423113), diluted 1:1000 in PBS (Thermo Scientific, 14190144). Cells were washed with PBS, stained with viability dye and relevant markers for 15 minutes, then washed with 1 mL FACS buffer (49 mL PBS + 100 µL 100 mM EDTA + 1 mL heat-inactivated FBS), fixed in 4% paraformaldehyde (ChemCruz, sc-281692), and resuspended in 200 µL FACS buffer. Data were acquired on BD Fortessa or Celesta instruments and analyzed using FlowJo v10.10.0. A minimum of 10,000 live (Zombie Violet-negative) events were collected per sample when possible.

### CRISPR/Cas9 gene knockouts

Protospacer targeting CRISPR RNA (crRNA) sequences for TAF1 and BRD1 were pre-designed by IDT or Broad Institute CRISPick (41, 42) and the sequences for Tat (CCUUAGGCAUCUCCUAUGGC) and non-targeting controls (ACGGAGGCUAAGCGUCGCAA) came from previous literature (34). crRNAs were synthesized by Integrated DNA Technologies (IDT) (Coralville, IA). CRISPR/Cas9 sgRNAs were made by mixing the crRNA and tracrRNA (IDT, 1072533) 1:1 at 100 µM, then annealing them in duplex buffer by heating to 95°C and letting them cool gradually to room temp using a thermocycler. RNPs were freshly assembled by combining 0.8 µL of Cas9 enzyme (IDT, 1081058), 0.8 µL of poly-glutamic acid (15 kDa, 100 mg/ mL) (Alamanda Polymers, CAS# 26247-79-0), and 1 µL of the annealed sgRNA (100 pmol), for a final molar ratio of 2:1 RNA to Cas9. To target TAF1 and BRD1, we multiplexed 3 single guide RNAs to improve knockout efficiency: TAF1 (1-UCACAGGGCACCGUCACGCG, 2-GGAGCGCCGGUACGUGCGC, 3-CGGUGUGGCCACUUAUCCUC), BRD1 (1-GUGGAGCUCGCGGCGUAUCG, 2-GGGGAAUUAUUCGGCGCGUA, 3-GGGCUCGUCGUAUAGUAGCG). We pelleted 1 million cells per nucleofection of the 2D10 cell line (Generous gift from Jonathan Karn) (43) at 90 RCF for 10 min. Cells were washed with PBS and pelleted at 90 RCF for 10 min, then resuspended in 20 µL of Lonza SE nucleofection buffer (Lonza, V4SC-1096). Cells were added to the RNPs and nucleofected using the CL-120 program on the Lonza 4D system. Immediately after nucleofection, cells were transferred into pre-warmed RPMI, rested at 37°C for 10 min, and then resuspended at 400,000 cells/mL in complete RPMI (34). Viral gene expression was measured via changes in GFP by flow cytometry at 5- and 7-days post nucleofection (DPN). At 5- and 7-DPN we collected 1 million cells for western blot to determine depletion efficacy. The distinct LRAs used to treat TAF1 depleted cells were vorinostat (500 nM, SelleckChem, S1047), prostratin (125 nM SigmaAldrich, P0077), or iBET-151 (125 nM, SelleckChem, S2780).

### Cell lysate and Western blotting

To generate protein lysates, approximately 1 million cells per sample were washed in PBS and lysed in RIPA buffer (Thermo Scientific, 89900) supplemented with complete protease inhibitors (Roche, 11697498001) and DNase I (Pierce, 88700) for 30 minutes at 4 °C with gentle agitation. Lysates were centrifuged at 16,000 × g for 10 minutes to remove debris, and protein concentrations were quantified using the DC Protein Assay (Bio-Rad, 5000111). Equal amounts of protein (5–10 µg per sample) were resolved on a 4–8% Tris-Acetate SDS-PAGE gel (ThermoFisher, EA03755BOX) and transferred onto a PVDF membrane (Invitrogen, IB23002). Membranes were blocked in TBS with 5% milk, incubated overnight with primary antibodies, and washed in TBST (TBS + 0.1% Tween-20). The primary antibodies used were anti-TAF1 (ab264327), anti-BRD1 (EPR12960), and anti-beta-actin (ab49900) as a loading control. HRP-conjugated secondary antibodies (Novex, A16035) were applied, and signal was developed using SuperSignal West Pico PLUS chemiluminescent substrate (Thermo Scientific, 34577) and imaged with a ChemiDoc MP system (Bio-Rad, Universal Hood III).

### Bulk RNA-sequencing

RNA was extracted from non-targeting control and TAF1-depleted 2D10 cells using the RNeasy Plus Kit (Qiagen, 74134) and eluted in elution buffer. RNA concentration was measured with the Qubit RNA HS Assay (Invitrogen), and integrity was assessed using a BioAnalyzer RNA Nano Kit on the TapeStation 4150 (Agilent). Libraries were prepared using the Watchmaker RNA Library Prep Kit (Techtum, 7K0077-096), quantified by Qubit, and sequenced on an Illumina NextSeq 2000 with 2×50 bp paired-end reads. Transcript counts were generated from aligned reads using RNASTAR (v2.7.1a), and differential expression analysis was performed with DESeq2 (v1.44.0).

### CUT&RUN

CUT&RUN was performed using the CUTANA CUT&RUN Kit (EpiCypher, 14-1048) with minor modifications. 2D10 cells were nucleofected with CRISPR/Cas9 RNPs targeting TAF1 or non-targeting controls as described, and collected in triplicate per antibody condition. For each replicate, 2 × 10⁶ cells were washed, bound to ConA-coated beads for 30 min at 4 °C, then divided into four groups and incubated overnight at 4 °C in antibody buffer containing 0.01% digitonin and 0.5 µg of antibody. Antibodies included: IgG (EpiCypher, 13-0042k), H3K4me3 (EpiCypher, 13-0060k), H3K9ac (Cell Signaling, 9649S), and RNA Polymerase II (Cell Signaling, 2629). DNA was quantified with the Qubit High Sensitivity dsDNA Assay (Invitrogen, Q328554).

Libraries were prepared using the CUTANA DNA Library Prep Kit (EpiCypher, 14-1001, Primer Set 1) with 14 cycles of PCR using dual-indexed barcodes. Libraries were quantified with Qubit and TapeStation 4150, pooled, and sequenced (50 bp paired-end, 20 million read pairs per sample) on an Illumina NextSeq 2000. Reads were demultiplexed with mkfastq, quality-checked with FastQC (v0.12.1), and trimmed with BBMap (v39.19). Alignment was performed using Bowtie2 (v2.4.5) to a custom genome combining GRCh38, HIV reference, and E. coli K12 (as pseudo-chromosomes). BAM files were indexed and merged using Samtools (v1.21). BigWig files for visualization were generated with bamCoverage (DeepTools v3.5.4), and coverage across the HIV genome was quantified using bigWigAverageOverBed (UCSC-tools).

### Statistical analysis

To account for potential plate effects in the compound screen, we used the exact stratified Wilcoxon test. This nonparametric method ranks observations within each plate and combines ranks across plates, enabling group comparisons while controlling for intra-plate variability. For grouped comparisons across experimental conditions, one-way ANOVA was applied. Dunnett’s multiple comparisons test was used when comparing multiple groups to a single control (e.g., non-targeting), while Tukey’s or Holm–Šidák corrections were used for all pairwise or selected group comparisons, as appropriate. These approaches provided robust statistical assessment across different replicate structures and experimental designs.

## Acknowledgements

This work was supported by the following grants from the National institutes of Health: NIAID #5-R01AI143381, NIAID #5-UM1AI164567. The funders had no role in study design, data collection and analysis, decision to publish, or preparation of the manuscript. Thanks to the UNC CFAR biostatistics core for the help in the statistical analysis.

---

**Table S1:**
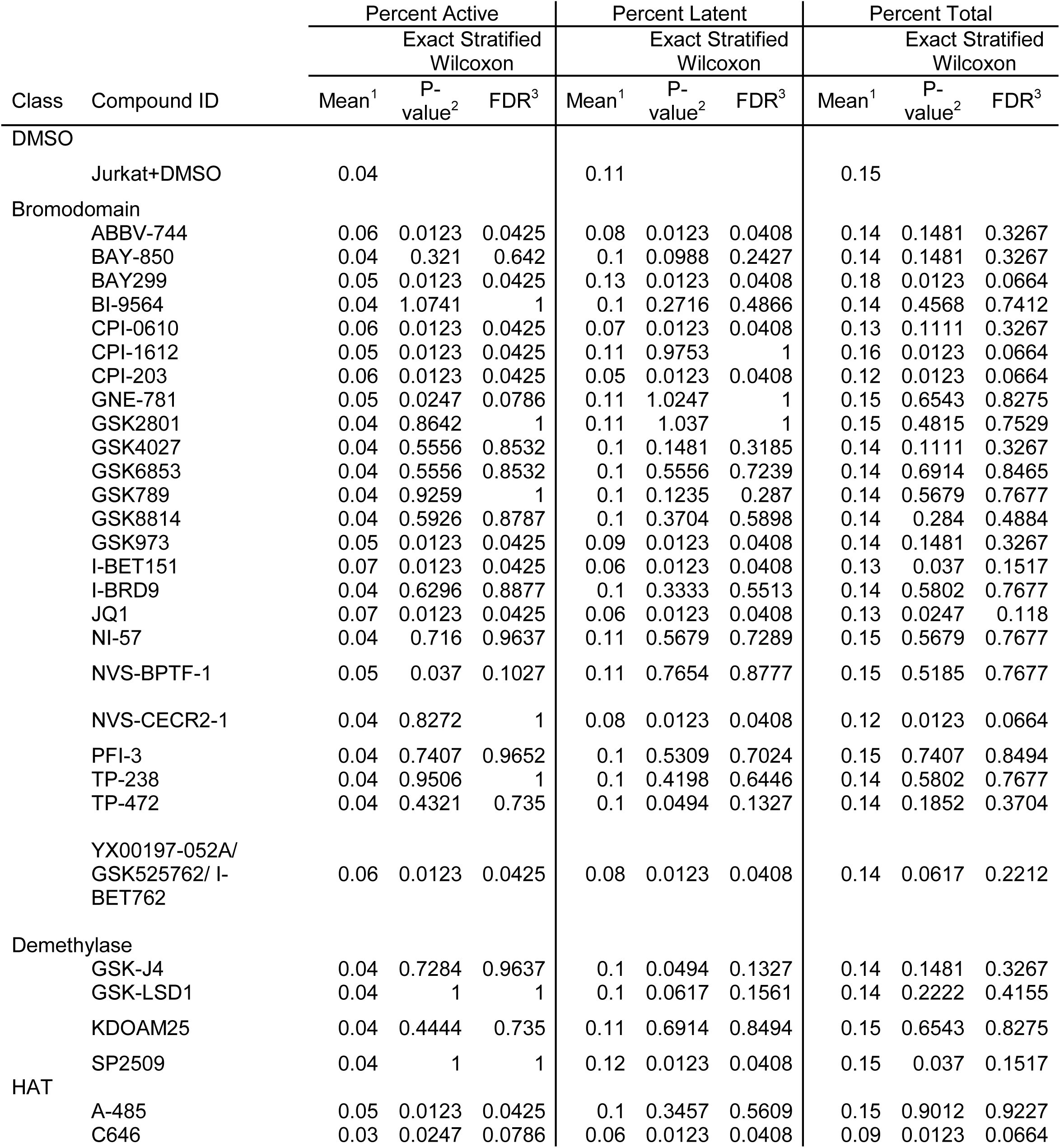
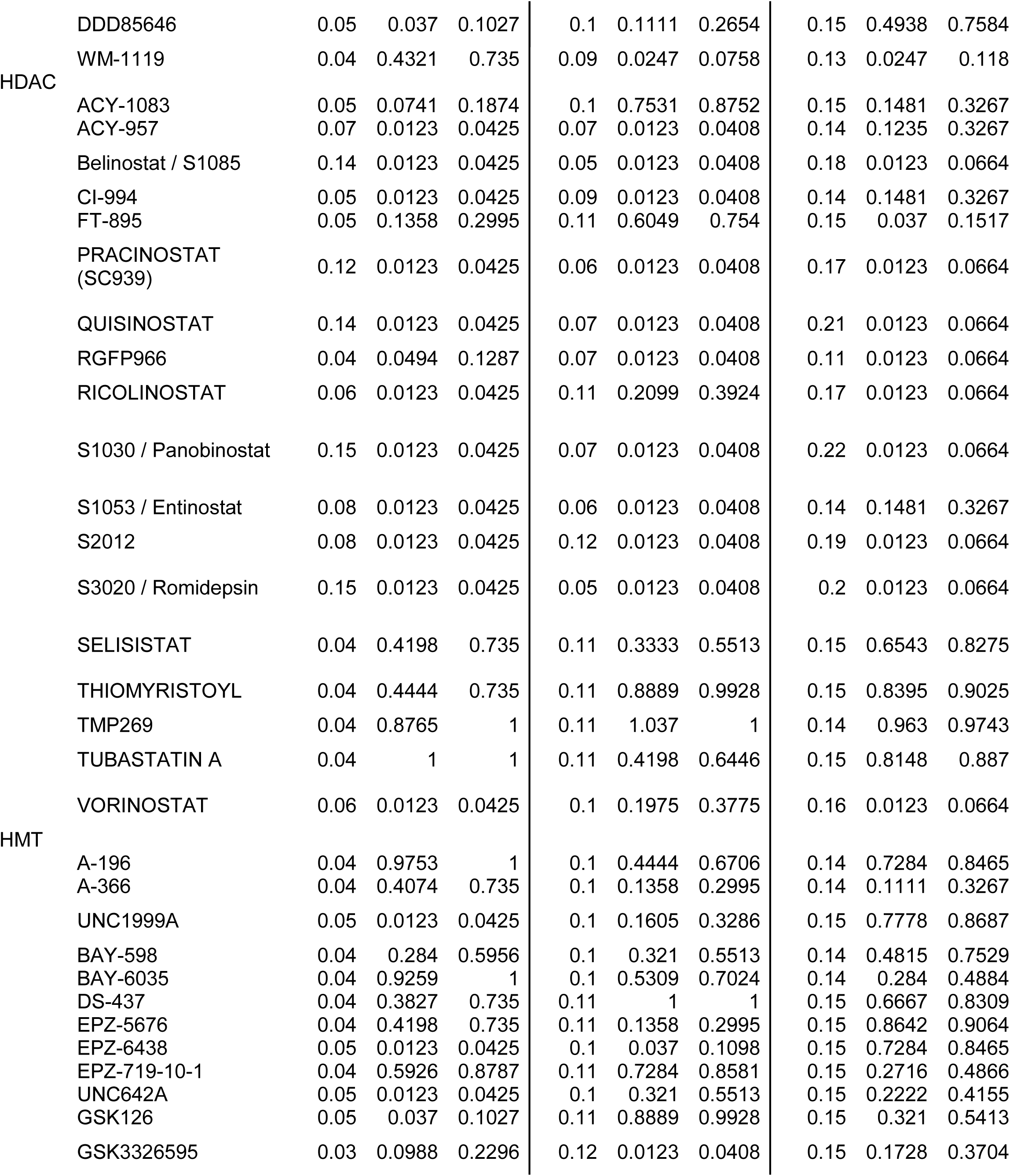
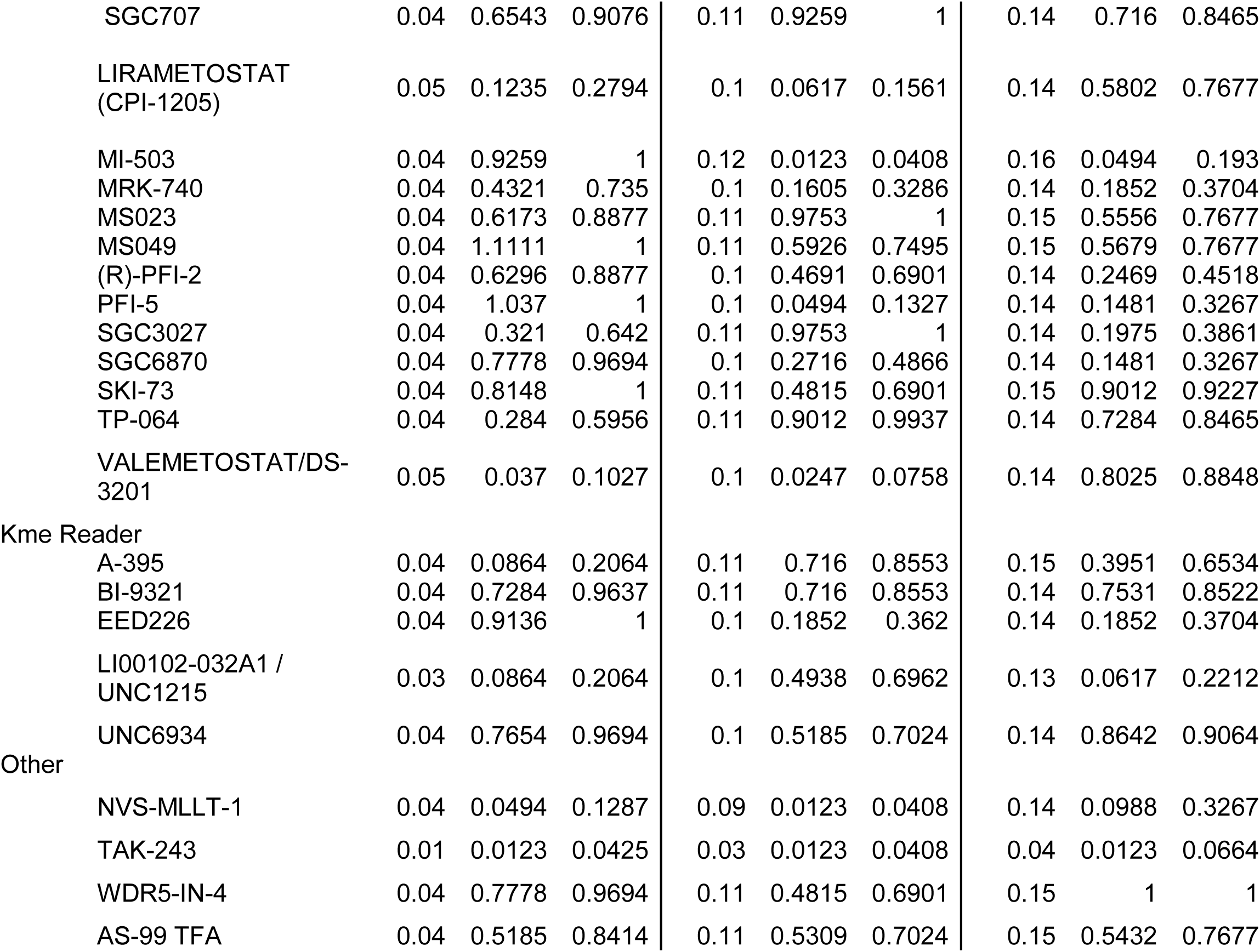
Raw dataset for compound screen. Raw data regarding abundance of total infected, actively infected and latently infected are shown for each compound included in the screen. Datapoints represent a fraction of overall cells and are an average of 4 technical replicates. Uncorrected and FDR corrected P values are shown.

## References

1. Pellowski JA, Price DM, Harrison AD, Tuthill EL, Myer L, Operario D, Lurie MN. 2019. A Systematic Review and Meta-analysis of Antiretroviral Therapy (ART) Adherence Interventions for Women Living with HIV. AIDS Behav 23:1998–2013.

2. Damulak PP, Ismail S, Abdul Manaf R, Mohd Said S, Agbaji O. 2021. Interventions to Improve Adherence to Antiretroviral Therapy (ART) in Sub-Saharan Africa: An Updated Systematic Review. 5. International Journal of Environmental Research and Public Health 18:2477.

3. Mateo-Urdiales A, Johnson S, Smith R, Nachega JB, Eshun-Wilson I. 2019. Rapid initiation of antiretroviral therapy for people living with HIV. Cochrane Database of Systematic Reviews 10.1002/14651858.CD012962.pub2.

4. Siliciano JD, Siliciano RF. 2022. In Vivo Dynamics of the Latent Reservoir for HIV-1: New Insights and Implications for Cure. Annual Review of Pathology: Mechanisms of Disease 17:271–294.

5. Li JZ, Melberg M, Kittilson A, Abdel-Mohsen M, Li Y, Aga E, Bosch RJ, Wonderlich ER, Kinslow J, Giron LB, Germanio CD, Pilkinton M, MacLaren L, Keefer M, Fox L, Barr L, Acosta E, Ananworanich J, Coombs R, Mellors J, Deeks S, Gandhi RT, Busch M, Landay A, Macatangay B, Smith DM, Team for the ACTGAS. 2024. Predictors of HIV rebound differ by timing of antiretroviral therapy initiation. JCI Insight 9:e173864.

6. Elliott JH, Wightman F, Solomon A, Ghneim K, Ahlers J, Cameron MJ, Smith MZ, Spelman T, McMahon J, Velayudham P, Brown G, Roney J, Watson J, Prince MH, Hoy JF, Chomont N, Fromentin R, Procopio FA, Zeidan J, Palmer S, Odevall L, Johnstone RW, Martin BP, Sinclair E, Deeks SG, Hazuda DJ, Cameron PU, Sékaly R-P, Lewin SR. 2014. Activation of HIV Transcription with Short-Course Vorinostat in HIV-Infected Patients on Suppressive Antiretroviral Therapy. PLOS Pathogens 10:e1004473.

7. Archin NM, Liberty AL, Kashuba AD, Choudhary SK, Kuruc JD, Crooks AM, Parker DC, Anderson EM, Kearney MF, Strain MC, Richman DD, Hudgens MG, Bosch RJ, Coffin JM, Eron JJ, Hazuda DJ, Margolis DM. 2012. Administration of vorinostat disrupts HIV-1 latency in patients on antiretroviral therapy. Nature 487:482–485.

8. Gutiérrez C, Serrano-Villar S, Madrid-Elena N, Pérez-Elías MJ, Martín ME, Barbas C, Ruipérez J, Muñoz E, Muñoz-Fernández MA, Castor T, Moreno S. 2016. Bryostatin-1 for latent virus reactivation in HIV-infected patients on antiretroviral therapy. AIDS 30:1385.

9. Chaillon A, Gianella S, Dellicour S, Rawlings SA, Schlub TE, Oliveira MFD, Ignacio C, Porrachia M, Vrancken B, Smith DM. 2020. HIV persists throughout deep tissues with repopulation from multiple anatomical sources. J Clin Invest 130:1699– 1712.

10. Archin NM, Kirchherr JL, Sung JA, Clutton G, Sholtis K, Xu Y, Allard B, Stuelke E, Kashuba AD, Kuruc JD, Eron J, Gay CL, Goonetilleke N, Margolis DM. 2017. Interval dosing with the HDAC inhibitor vorinostat effectively reverses HIV latency. 8. J Clin Invest 127:3126–3135.

11. Rasmussen TA, Tolstrup M, Brinkmann CR, Olesen R, Erikstrup C, Solomon A, Winckelmann A, Palmer S, Dinarello C, Buzon M, Lichterfeld M, Lewin SR, Østergaard L, Søgaard OS. 2014. Panobinostat, a histone deacetylase inhibitor, for latent-virus reactivation in HIV-infected patients on suppressive antiretroviral therapy: a phase 1/2, single group, clinical trial. The Lancet HIV 1:e13–e21.

12. Søgaard OS, Graversen ME, Leth S, Olesen R, Brinkmann CR, Nissen SK, Kjaer AS, Schleimann MH, Denton PW, Hey-Cunningham WJ, Koelsch KK, Pantaleo G, Krogsgaard K, Sommerfelt M, Fromentin R, Chomont N, Rasmussen TA, Østergaard L, Tolstrup M. 2015. The Depsipeptide Romidepsin Reverses HIV-1 Latency In Vivo. PLoS Pathog 11:e1005142.

13. Jones JE, Gunderson CE, Wigdahl B, Nonnemacher MR. 2025. Impact of chromatin on HIV-1 latency: a multi-dimensional perspective. Epigenetics & Chromatin 18:9.

14. Nixon CC, Mavigner M, Sampey GC, Brooks AD, Spagnuolo RA, Irlbeck DM, Mattingly C, Ho PT, Schoof N, Cammon CG, Tharp GK, Kanke M, Wang Z, Cleary RA, Upadhyay AA, De C, Wills SR, Falcinelli SD, Galardi C, Walum H, Schramm NJ, Deutsch J, Lifson JD, Fennessey CM, Keele BF, Jean S, Maguire S, Liao B, Browne EP, Ferris RG, Brehm JH, Favre D, Vanderford TH, Bosinger SE, Jones CD, Routy J-P, Archin NM, Margolis DM, Wahl A, Dunham RM, Silvestri G, Chahroudi A, Garcia JV. 2020. Systemic HIV and SIV latency reversal via non-canonical NF-κB signalling in vivo. Nature 578:160–165.

15. Jütte BB, Love L, Svensson JP. 2023. Molecular Mechanisms of HIV-1 Latency from a Chromatin and Epigenetic Perspective. Curr Clin Micro Rpt 10:246–254.

16. Castillo J, López-Rodas G, Franco L. 2017. Histone Post-Translational Modifications and Nucleosome Organisation in Transcriptional Regulation: Some Open Questions, p. 65–92. In Atassi, MZ (ed.), Protein Reviews: Volume 18. Springer, Singapore.

17. Letchumanan P, Theva Das K. 2025. The role of genetic diversity, epigenetic regulation, and sex-based differences in HIV cure research: a comprehensive review. Epigenetics & Chromatin 18:1.

18. Mori L, Valente ST. 2020. Key Players in HIV-1 Transcriptional Regulation: Targets for a Functional Cure. Viruses 12:529.

19. Verdikt R, Hernalsteens O, Van Lint C. 2021. Epigenetic Mechanisms of HIV-1 Persistence. 5. Vaccines 9:514.

20. Margolis DM, Archin NM, Cohen MS, Eron JJ, Ferrari G, Garcia JV, Gay CL, Goonetilleke N, Joseph SB, Swanstrom R, Turner A-MW, Wahl A. 2020. Curing HIV: Seeking to Target and Clear Persistent Infection. 1. Cell 181:189–206.

21. Cillo AR, Sobolewski MD, Bosch RJ, Fyne E, Piatak M, Coffin JM, Mellors JW. 2014. Quantification of HIV-1 latency reversal in resting CD4+ T cells from patients on suppressive antiretroviral therapy. Proc Natl Acad Sci U S A 111:7078–7083.

22. Bouché L, Christ CD, Siegel S, Fernández-Montalván AE, Holton SJ, Fedorov O, Ter Laak A, Sugawara T, Stöckigt D, Tallant C, Bennett J, Monteiro O, Díaz-Sáez L, Siejka P, Meier J, Pütter V, Weiske J, Müller S, Huber KVM, Hartung IV, Haendler B. 2017. Benzoisoquinolinediones as Potent and Selective Inhibitors of BRPF2 and TAF1/TAF1L Bromodomains. J Med Chem 60:4002–4022.

23. Battivelli E, Dahabieh MS, Abdel-Mohsen M, Svensson JP, Tojal Da Silva I, Cohn LB, Gramatica A, Deeks S, Greene WC, Pillai SK, Verdin E. 2018. Distinct chromatin functional states correlate with HIV latency reactivation in infected primary CD4+ T cells. Elife 7:e34655.

24. Pearson R, Kim YK, Hokello J, Lassen K, Friedman J, Tyagi M, Karn J. 2008. Epigenetic Silencing of Human Immunodeficiency Virus (HIV) Transcription by Formation of Restrictive Chromatin Structures at the Viral Long Terminal Repeat Drives the Progressive Entry of HIV into Latency. Journal of Virology 82:12291– 12303.

25. Bradley T, Ferrari G, Haynes BF, Margolis DM, Browne EP. 2018. Single-Cell Analysis of Quiescent HIV Infection Reveals Host Transcriptional Profiles that Regulate Proviral Latency. Cell Rep 25:107–117.e3.

26. Raha T, Cheng SWG, Green MR. 2005. HIV-1 Tat Stimulates Transcription Complex Assembly through Recruitment of TBP in the Absence of TAFs. PLOS Biology 3:e44.

27. Bernardini A, Mukherjee P, Scheer E, Kamenova I, Antonova S, Mendoza Sanchez PK, Yayli G, Morlet B, Timmers HTM, Tora L. 2023. Hierarchical TAF1-dependent co-translational assembly of the basal transcription factor TFIID. Nat Struct Mol Biol 30:1141–1152.

28. Patel AB, Greber BJ, Nogales E. 2020. Recent insights into the structure of TFIID, its assembly, and its binding to core promoter. Current Opinion in Structural Biology 61:17–24.

29. Ashokkumar M, Hafer TL, Felton A, Archin NM, Margolis DM, Emerman M, Browne EP. 2025. A targeted CRISPR screen identifies ETS1 as a regulator of HIV-1 latency. PLOS Pathogens 21:e1012467.

30. Chou TC, Maggirwar NS, Marsden MD. 2024. HIV Persistence, Latency, and Cure Approaches: Where Are We Now? 7. Viruses 16:1163.

31. Kreider EF, Bar KJ. 2022. HIV-1 Reservoir Persistence and Decay: Implications for Cure Strategies. Curr HIV/AIDS Rep 19:194–206.

32. Siliciano JD, Siliciano RF. 2024. HIV cure: The daunting scale of the problem. Science 383:703–705.

33. Falcinelli SD, Peterson JJ, Turner A-MW, Irlbeck D, Read J, Raines SLM, James KS, Sutton C, Sanchez A, Emery A, Sampey G, Ferris R, Allard B, Ghofrani S, Kirchherr JL, Baker C, Kuruc JD, Gay CL, James LI, Wu G, Zuck P, Rioja I, Furze RC, Prinjha RK, Howell BJ, Swanstrom R, Browne EP, Strahl BD, Dunham RM, Archin NM, Margolis DM. Combined noncanonical NF-κB agonism and targeted BET bromodomain inhibition reverse HIV latency ex vivo. J Clin Invest 132:e157281.

34. Peterson JJ, Lewis CA, Burgos SD, Manickam A, Xu Y, Rowley AA, Clutton G, Richardson B, Zou F, Simon JM, Margolis DM, Goonetilleke N, Browne EP. 2023. A histone deacetylase network regulates epigenetic reprogramming and viral silencing in HIV-infected cells. Cell Chemical Biology 30:1617–1633.e9.

35. Wu G, Swanson M, Talla A, Graham D, Strizki J, Gorman D, Barnard RJ, Blair W, Søgaard OS, Tolstrup M, Østergaard L, Rasmussen TA, Sekaly R-P, Archin NM, Margolis DM, Hazuda DJ, Howell BJ. 2017. HDAC inhibition induces HIV-1 protein and enables immune-based clearance following latency reversal. 16. JCI Insight 2.

36. Gegonne A, Weissman JD, Singer DS. 2001. TAFII55 binding to TAFII250 inhibits its acetyltransferase activity. Proceedings of the National Academy of Sciences 98:12432–12437.

37. Weissman JD, Brown JA, Howcroft TK, Hwang J, Chawla A, Roche PA, Schiltz L, Nakatani Y, Singer DS. 1998. HIV-1 Tat binds TAFII250 and represses TAFII250-dependent transcription of major histocompatibility class I genes. Proceedings of the National Academy of Sciences 95:11601–11606.

38. Mzingwane ML, Tiemessen CT. 2017. Mechanisms of HIV persistence in HIV reservoirs. Reviews in Medical Virology 27:e1924.

39. Gálvez C, Grau-Expósito J, Urrea V, Clotet B, Falcó V, Buzón MJ, Martinez-Picado J. 2021. Atlas of the HIV-1 Reservoir in Peripheral CD4 T Cells of Individuals on Successful Antiretroviral Therapy. mBio 12:e03078–21.

40. Nixon CC, Mavigner M, Sampey GC, Brooks AD, Spagnuolo RA, Irlbeck DM, Mattingly C, Ho PT, Schoof N, Cammon CG, Tharp GK, Kanke M, Wang Z, Cleary RA, Upadhyay AA, De C, Wills SR, Falcinelli SD, Galardi C, Walum H, Schramm NJ, Deutsch J, Lifson JD, Fennessey CM, Keele BF, Jean S, Maguire S, Liao B, Browne EP, Ferris RG, Brehm JH, Favre D, Vanderford TH, Bosinger SE, Jones CD, Routy J-P, Archin NM, Margolis DM, Wahl A, Dunham RM, Silvestri G, Chahroudi A, Garcia JV. 2020. Systemic HIV and SIV latency reversal via non-canonical NF-κB signalling in vivo. Nature 578:160–165.

41. Kim HK, Min S, Song M, Jung S, Choi JW, Kim Y, Lee S, Yoon S, Kim H (Henry). 2018. Deep learning improves prediction of CRISPR–Cpf1 guide RNA activity. Nat Biotechnol 36:239–241.

42. DeWeirdt PC, Sanson KR, Sangree AK, Hegde M, Hanna RE, Feeley MN, Griffith AL, Teng T, Borys SM, Strand C, Joung JK, Kleinstiver BP, Pan X, Huang A, Doench JG. 2021. Optimization of AsCas12a for combinatorial genetic screens in human cells. Nat Biotechnol 39:94–104.

43. Pearson R, Kim YK, Hokello J, Lassen K, Friedman J, Tyagi M, Karn J. 2008. Epigenetic Silencing of Human Immunodeficiency Virus (HIV) Transcription by Formation of Restrictive Chromatin Structures at the Viral Long Terminal Repeat Drives the Progressive Entry of HIV into Latency. Journal of Virology 82:12291– 12303.

